# Natural Oil Blend Formulation (NOBF) as an anti- African Swine Fever Virus (ASFV) agent in Primary Porcine Alveolar Macrophases (PAM) cells of Swine

**DOI:** 10.1101/2020.10.09.332890

**Authors:** Haig Yousef Babikian, Rajeev Kumar Jha, Quang Lam Truong, Lan Thi Nguyen, Hoa Thi Nguyen, Thanh Long To

## Abstract

African swine fever is one of the severe pathogens of swine. It has a significant impact on production and on economics. So far, there are no known remedies, such as vaccines or drugs, reported. The natural oil blend formulation (NOBF) successfully tested against the African swine fever virus (ASFV) in *in vitro* conditions. The natural oil blend formulation (NOBF) combines *Eucalyptus globulus*, *Pinus sylvestris*, and *Lavandula latifolia*. We used a two-fold serial dilution to test the NOBF formulation dose. The *in vitro* trial results demonstrated that NOBF up to dilution 13 or 0.000625 ml deactivates the lethal dose 105HAD50 of ASFV. There was no hemadsorption (Rosetta formation) up to dilution 12 or 0.00125 ml of NOBF. The Ct value of the NOBF group at 96 hours post-infection was the same as the initial value or lower (25), whereas the Ct value of positive controls increased several folds (17.84). The *in vitro* trial demonstrated that NOBF can deactivate the African swine fever virus.

**HIGHLIGHTS:**

- The natural oil blend formulation (NOBF) was formulated using three natural oils, i.e., *Eucalyptus globulus*, *Pinus sylvestris*, and *Lavandula latifolia*.
- The in vitro trial was conducted using porcine alveolar macrophages (PAMs) and further passaged in the PAMs; the stock used in the present study was that obtained after the 15th passage.
- The natural oil blend formulation (NOBF) showed protection against ASF virus up to a dilution of 13 or 0.000625 of a dilution of 16 or 0.000078 ml that was tried.
- The real-time PCR analysis showed that the virus did not replicate in the NOBF group, which implies that either ASFV growth was inhibited in the presence of NOBF or that it was inactivated.
- The *in vitro* trial outcome showed that NOBF has anti-ASFV properties.

## INTRODUCTION

African Swine Fever Virus (ASFV), reported deadly for pigs. It is listed as a “notifiable disease” by the OIE due to high illness rates and a high mortality rate, up to 100%, and substantial financial losses (Penrith et al., 2013 and Rowlands et al. 2008, Halasa et al. 2016). Further spread of ASF to China has had disastrous consequences, especially in lieu of the fact that China contains more than half of the world’s pig population (Sánchez-Cordón et al., 2018). To date, as far as Vietnam is concerned, ASF has appeared in all 63 provinces of Vietnam, has destroyed more than 5.6 million pigs (more than 20% of total pigs), has decreased pork production by 8.3%, and has affected mainly small-scale farms (WOAH 2019).

The typical signs and symptoms of ASF are high fever, decreased appetite and weakness, difficulty in standing, red or blue blotches on the skin (particularly around ears and snout), and especially in sows, the symptoms of miscarriage, stillbirths, and weak litters can occur (WOAH 2019). The symptoms, like, diarrhea, vomiting, and difficulty in breathing or coughing, can also occur with the disease (WOAH 2019).

African swine fever virus is a large, enveloped, and structurally complex DNA virus with the *Asfarviridae* family’s icosahedral morphology.

The virus can persist for a long time in the environment, carcasses, and various swine products. The vectors and carriers of the ASF virus are warthogs (*Phacochoerus africanus*), bushpigs (*Potamochoerus porcus*, *P. larvatus*), and soft ticks (*Ornithodoros moubata*) (Sánchez-Cordón et al., 2018) in which the virus is transmitted trans-staidly and through transovarial routes (Hess et al., 1987).

The role of natural oils as antiviral components is well known. In both crude and pure forms, as a standardized compound, natural products are significant components with antiviral properties (Jassim and Naji, 2003). A formulation was developed by blending three natural oils, *Eucalyptus globulus*, *Pinus sylvestris*, and *Lavandula latifolia*, with antiviral properties. Cineole, the significant component of eucalyptus oil, has potent anti-inflammatory and anti-microbial properties (Jun et al. 2013). Cineole is also known to treat the respiratory tract’s primary viral infections (Muller et al. 2016 and Li et al. 2016). Linalool, a major component of lavender oil, has shown antiviral activities (Choi 2018 and Takizawa et al. 2001). Isobornyl acetate extracted from pine oil has anti-microbial properties (Li et al. 2016).

In the present study, the efficacy of the natural oil bled formulation was evaluated against African swine fever using porcine alveolar macrophage (PAM) cells of swine in an *in vitro* medium.

## MATERIALS AND METHODS

### 1. Trial Station

In this trial, we selected 7-10 weeks-old healthy pigs negative for ASFV, PCV2, CSFV, PRRSV as well as negative for ASFV Ab. The animals were housed and used in isolated area in the Biosecurity Animal Facility center of the Vietnam National University of Agriculture (VNUA), Hanoi, Vietnam. The Ministry of Agriculture and Rural Development approved by the VNUA Animal Care and Use Committee granted permission for this trial. The study period was between October 2019 and January 2020.

### 2. Natural oil blend Formulation (NOBF)

A formulation was developed by mixing an essential oil blend using three oils, *Eucalyptus globulus*, *Pinus sylvestris*, and *Lavandula latifolia*, in a determined concentration. The formulation was pre-tested on animals for toxicity and tolerance level. We followed a two-fold dilution procedure to obtain the optimum dose of the application. The natural oil blend formulation (NOBF) was serially diluted from 2.5 ml to 0.000078 ml (dilution 1 to dilution 16) to perform the *in vitro* trials.

### 3. Toxicity Test of Natural oil blend Formulation (NOBF)

The toxicity level of NOBF was tested at two levels, first level test was performed in the PAM cells whereas the tolerance level of NOBF was tested in live pigs. The NOBF in 16 double-fold dilutions of 2.5 ml to 0.000078 ml were mixed with the PAM cells and the ASFV. The cell viability was tested under a microscope at every 24-hour interval.

### 4. Active ingredient identification in NOBF

The gas chromatography technique was used to extract and identify the active ingredients with anti-ASFV properties. Gas Agilent 6890 and 7890 Gas Chromatography - Flame Ionization Detector (GC-FID) Shimadzu GC 2014) analysis was conducted at the Faculty of Pharmacy laboratory at the University of Indonesia. The details of the standards used in in this analysis are as follows:

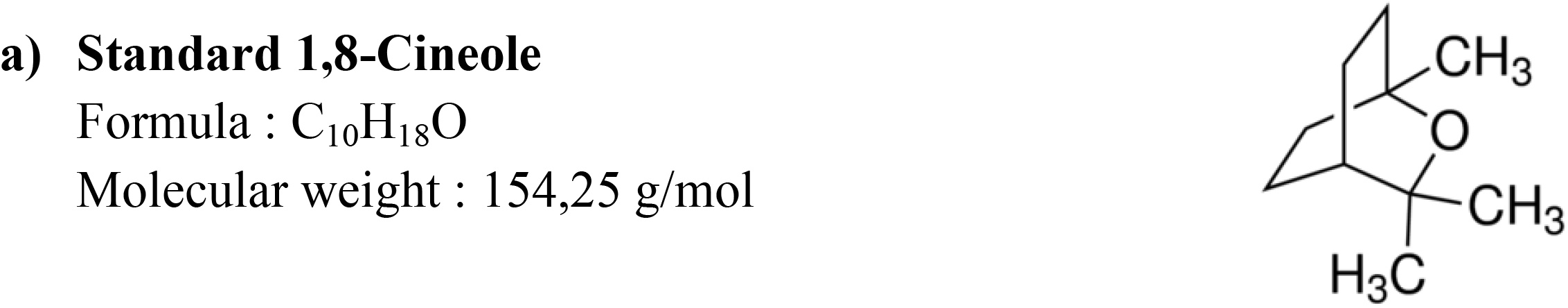
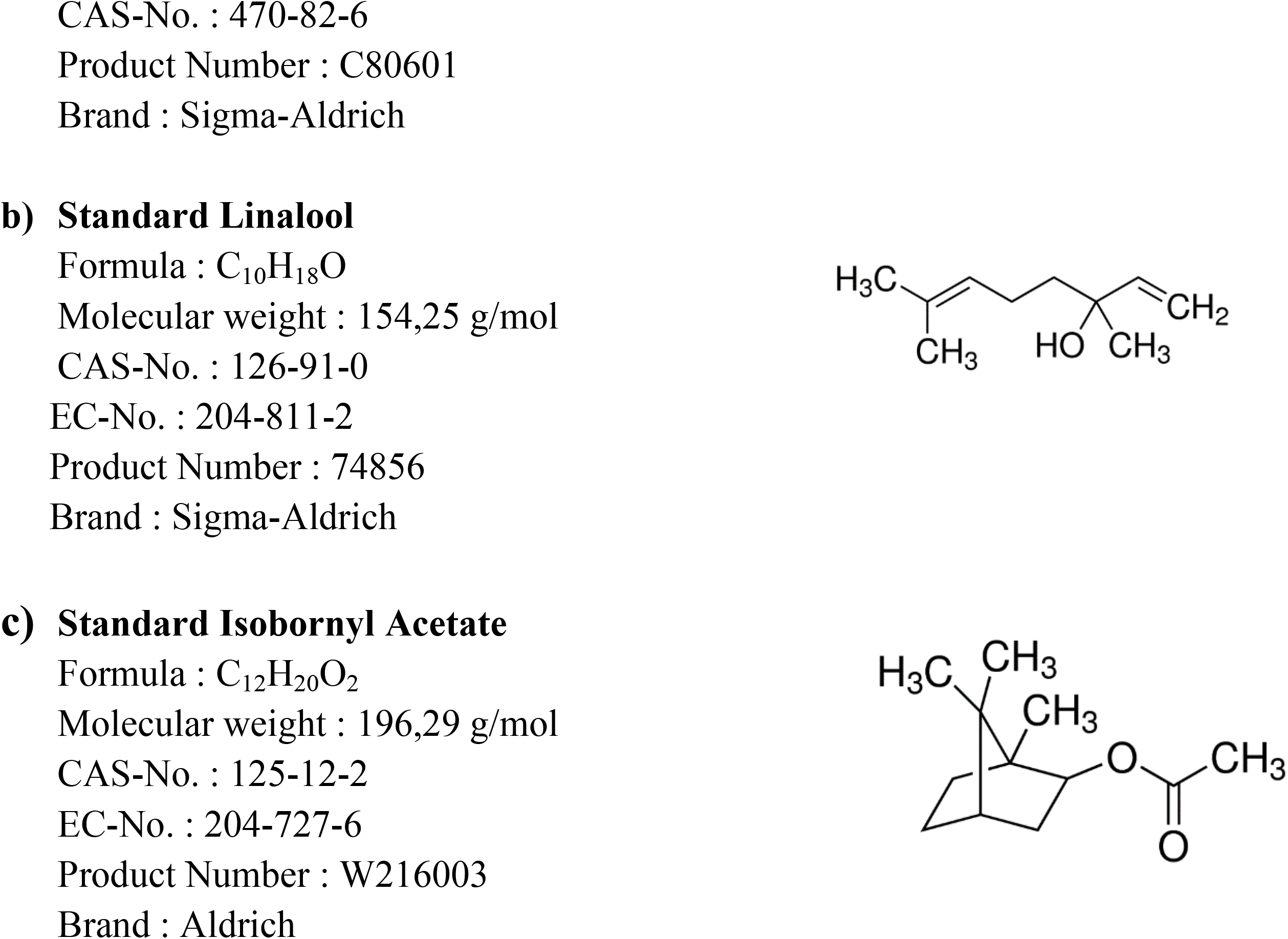

### 5. Primary porcine alveolar macrophages (PAM) cells

Our team collected the primary porcine alveolar macrophages (PAMs) from 7 weeks-old healthy pigs (negative for ASFV, PCV2, CSFV, PRRSV). We maintained the cells in the growth medium, including an RPMI 1640 medium (Gibco, USA) supplemented with 10% fatal calf serum (FCS; Gibco, USA), 1% penicillin-streptomycin solution (Gibco, USA) at 37°C with 5% CO_2_. We also prepared red blood cells from EDTA-treated swine blood by using Percoll (GE Healthcare, USA) and kept it in an RPMI 1640 medium (Gibco, USA), 1% penicillin-streptomycin solution, and maintained it at 4°C until use.

### 6. African swine fever virus preparation

The VNUA-ASFV-L01/HN/04/19 virus strain was isolated from a pig with apparent symptoms in Thai Binh province, Vietnam. The virus strain was purified and quantified in the molecular biology laboratory of VNUA, Vietnam. The lethal and sublethal doses were optimized in the controlled conditions. The ASFV strain VNUA01/04.2019 was adapted to grow in porcine alveolar macrophages (PAMs), and further passaged in PAMs. The stock used in the present study was that obtained after the 15^th^ passage. Briefly, the PAMs were infected at a multiplicity of infection (MOI) of 0.1 with VNUA-ASFV-L01/HN/04/19 in the growth medium, including an RPMI 1640 medium supplemented with 10% fatal calf serum (FCS) and 1% penicillin-streptomycin solution. We added to each well the maintenance medium containing an RPMI 1640 medium supplemented with 5% fatal calf serum (FCS), 1% penicillin-streptomycin solution, and 2% suspension of red blood cells. The team performed the ASFV titration on the PAMs cultures in 96-well plates. The presence of ASFV was assessed by hemadsorption (HAD). The observation of HAD was for 5 days, and the calculation of 50% HAD observation (HAD50) was performed by using the method described by Reed and Muench (1938).

### 7. Anti-viral activity of natural oil blend formulation (NOBF) using an in vitro medium

The PAMs were grown on 48-well tissue culture plates using a growth medium, including an RPMI 1640 medium supplemented with 10% fatal calf serum (FCS) and 1% penicillin-streptomycin solution. The natural oil blend formulation (NOBF) was serially two-fold diluted (Table 1) in a warmed RPMI 1640 to prepare the working stocks after 3 hours of incubation of serially 2-fold-diluted NOBF with VNUA-ASFV-L01/HN/05/19.

**Table 1:**
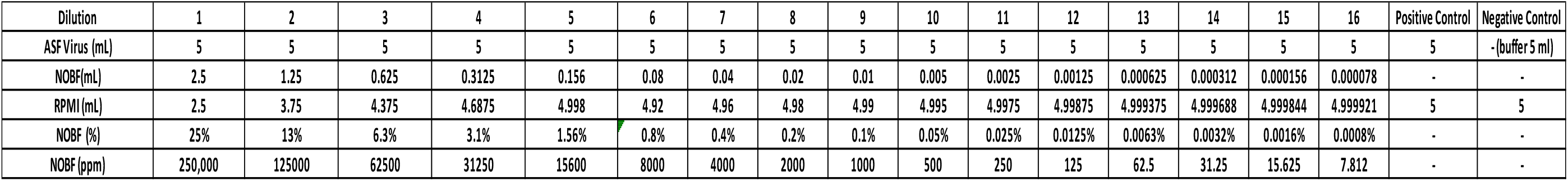
Natural oil blend formulation (NOBF) mixed with RPMI and ASFV in serial fold 2 dilution. The 5 ml of VNUA-ASFV-L01/HN/05/19 mixed in each tube except negative control. The NOBF was serially 2-fold diluted up to 7.8 ppm.

| Dilution | 1 | 2 | 3 | 4 | 5 | 6 | 7 | 8 | 9 | 10 | 11 | 12 | 13 | 14 | 15 | 16 | Positive Control | Negative Control |
| --- | --- | --- | --- | --- | --- | --- | --- | --- | --- | --- | --- | --- | --- | --- | --- | --- | --- | --- |
| ASF Virus (mL) | 5 | 5 | 5 | 5 | 5 | 5 | 5 | 5 | 5 | 5 | 5 | 5 | 5 | 5 | 5 | 5 | 5 | -(buffer 5 ml) |
| NOBF (mL) | 2.5 | 1.25 | 0.625 | 0.3125 | 0.156 | 0.08 | 0.04 | 0.02 | 0.01 | 0.005 | 0.0025 | 0.00125 | 0.000625 | 0.000312 | 0.000156 | 0.000078 | - | - |
| RPMI (mL) | 2.5 | 3.75 | 4.375 | 4.6875 | 4.998 | 4.92 | 4.96 | 4.98 | 4.99 | 4.995 | 4.9975 | 4.99875 | 4.999375 | 4.999688 | 4.999844 | 4.999921 | 5 | 5 |
| NOBF (%) | 25% | 13% | 6.3% | 3.1% | 1.56% | 0.8% | 0.4% | 0.2% | 0.1% | 0.05% | 0.025% | 0.0125% | 0.0063% | 0.0032% | 0.0016% | 0.0008% | - | - |
| NOBF (ppm) | 250,000 | 125000 | 62500 | 31250 | 15600 | 8000 | 4000 | 2000 | 1000 | 500 | 250 | 125 | 62.5 | 31.25 | 15.625 | 7.812 | - | - |

At the 10^5^HAD50 at the ratio of 1:1, duplicate cultures were infected with the corresponding virus in a diluted volume of a medium containing the NOBF at 37°C in an atmosphere of 5% CO_2_ for 1 hour (Table 1 and Figure 1). Cultures were added to the maintenance medium and incubated until a massive cytopathic effect, such as hemadsorption (HAD) or rosette formation. Rosette formation was observed daily by an inverted microscope for 4-5 days. After four freeze-thaw cycles, the supernatants were assessed for the ASFV virus by real-time PCR using the recommendations in the OIE manual described by King et al. 2003.

**Figure 1:**
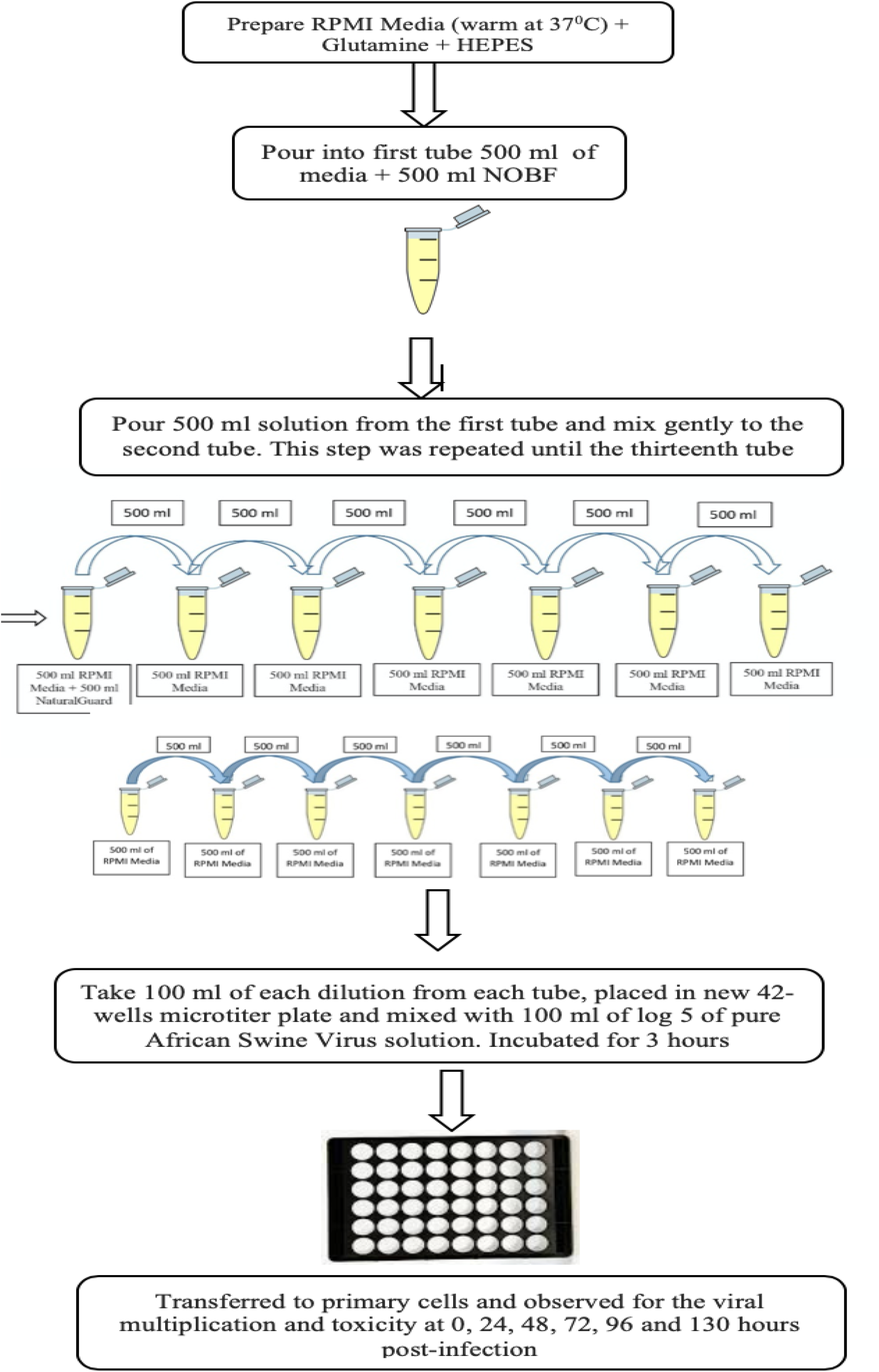
Diagrammatic illustration of in vitro trial steps preparation. RPMI media, NOBF and ASFV prepared and mixed using 2-fold dilution method. No virus was mixed in negative control. Microscopic observation was made at 0 h, 24 h, 48 h, 72 h, 96 h and 130 h intervals.

## RESULTS AND DISCUSSION

African swine fever is a highly contagious fatal acute hemorrhagic viral disease of pigs that currently has no treatment or vaccination protocol. It threatens the pig industry worldwide. For farmers, managing recent outbreaks of infectious viral diseases remains a significant worldwide problem, and there is a need to find substances with intracellular and extracellular anti-viral properties (Jurgen et al. 2009). Simoes et al. (2015) stated that ASFV precisely activates the Ataxia Telangiectasia Mutated Rad-3 related (ATR) pathway in ASFV-infected swine MDMs in the early phase of infection, most probably because the ASFV genome is recognized as foreign DNA. They also detected morphological changes of promyelocytic leukemia nuclear bodies (PML-NBs), nuclear speckles, and Cajal bodies found in ASFV-infected swine MDMS. This suggests the viral modulation of cellular anti-viral responses and cellular transcription. Simoes et al. (2019) demonstrated that *in vitro* inhibition of ASFV-topoisomerase II disrupts viral replications, contributing to natural strategies for vaccine candidate development.

### 1. The active ingredient natural oil blend formulation (NOBF)

The gas chromatography technique measures the active ingredients present in the blended oil. By comparing with chromatographs of pure standards, GC analysis identified the presence of anti-viral compounds, namely, A) Cineole B) Linalool and C) Isobornyl acetate in the NOBF (Figure 2). The result shows that Cineole was present at a retention time of 3.048 minutes with a relative percentage area of peak around 99.4%, which makes it the primary compound in the NOBF. The second compound identified is Linalool at a retention time of 5.966 minutes and a relative percentage area of peak around 0.31%. The minor compound identified is Isobornyl acetate at a retention time of 3.767 and a relative percentage area of peak around 0.28% (Table 2).

**Figure 2:**
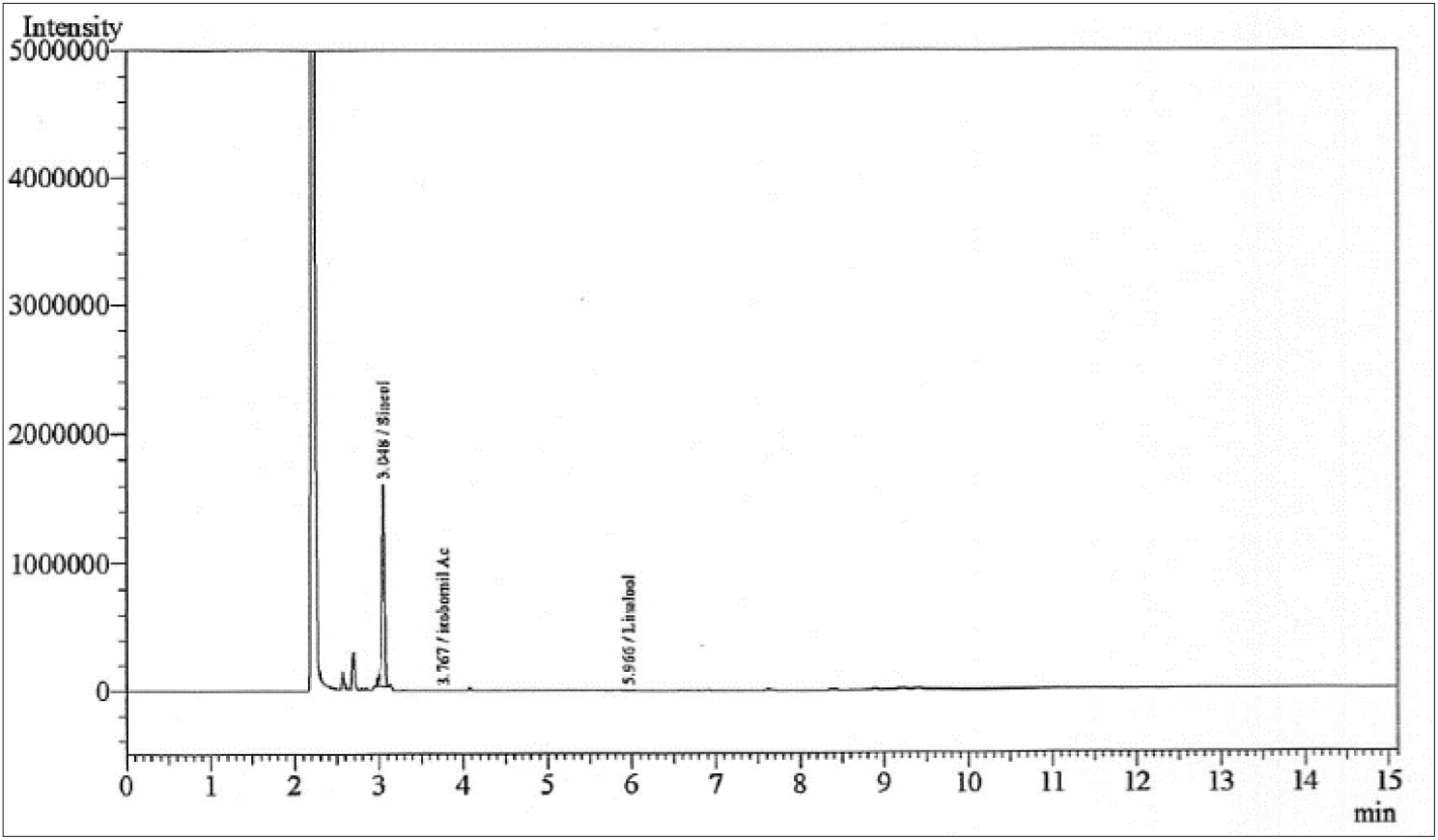
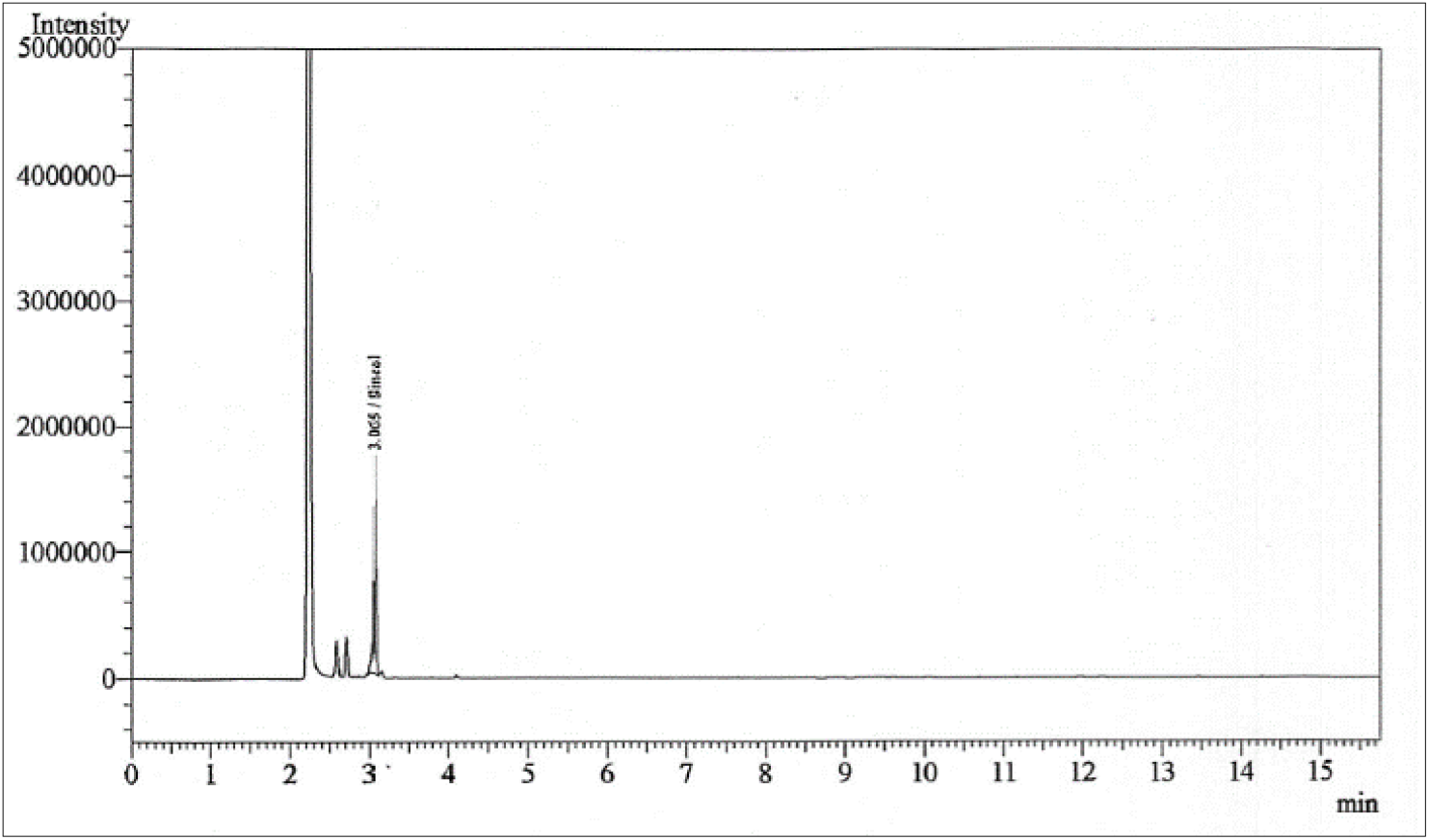
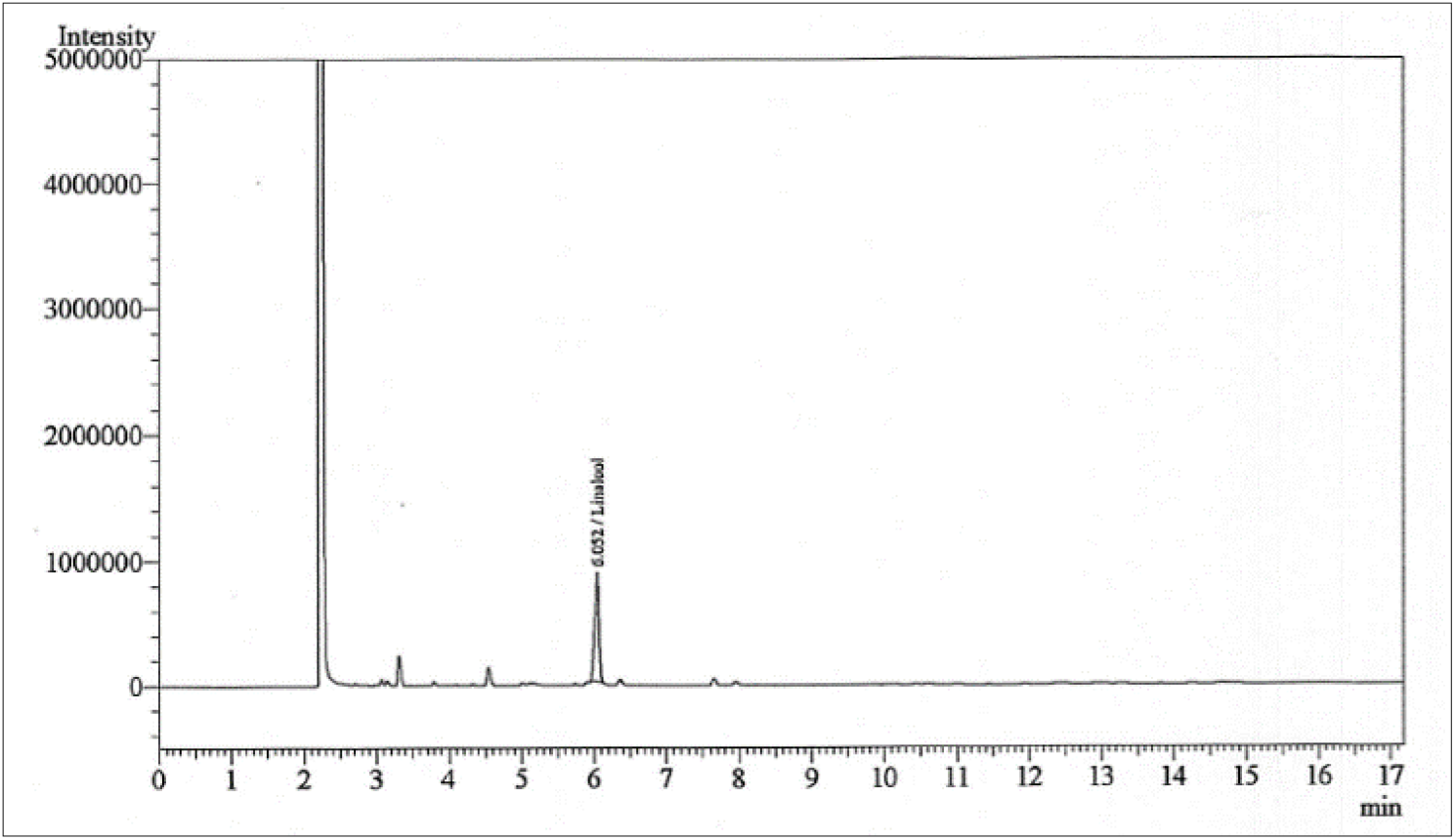
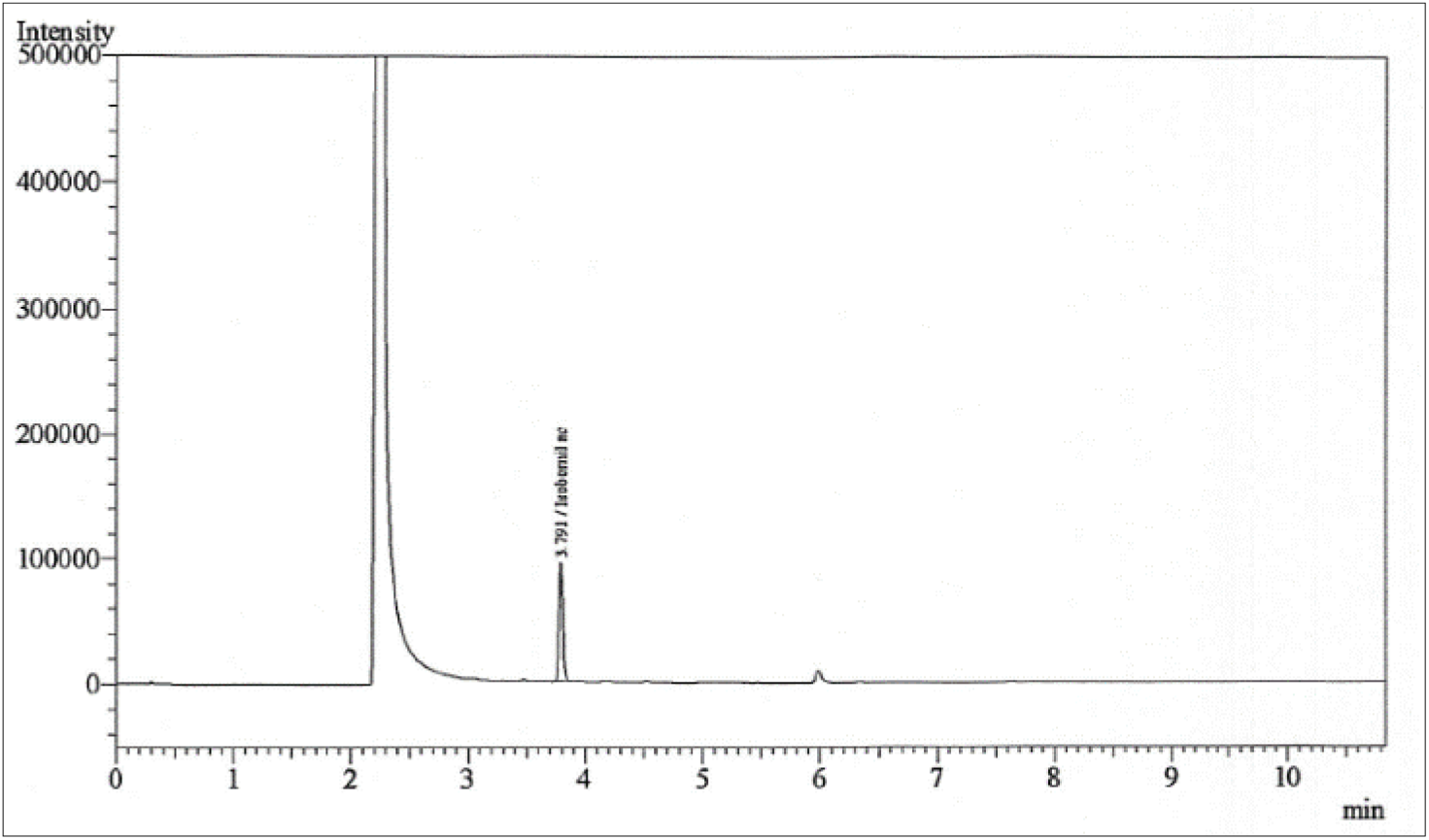
GC-FID Chromatogram of NOBF. a) Chromatogram of complete NOBF compound’s peak. b) Chromatogram of Cineole from NOBF c) Chromatogram of Linalool from NOBF and, d) Chromatogram of Isobornyl acetate from NOBF.

**Table 2 :** GC-FID Chromatogram of NOBF in tabular form with complete details of measurement and analysis.

| Peak No | Name | Ret. Time | Area | Height | HETP | Resolution | Tailing Factor | k' | Separation |
| --- | --- | --- | --- | --- | --- | --- | --- | --- | --- |
| 1 | Cineole | 3.048 | 3516085 | 11587085 | 0.000 | 0.000 | 0.831 | 0.000 | 0.000 |
| 2 | Linalool | 5.966 | 11108 | 4328 | 0.000 | 12.943 | 0.840 | 0.236 | 0.000 |
| 3 | Isobornyl acetate | 3.767 | 10037 | 2854 | 0.000 | 30.333 | 0.000 | 0.957 | 4.057 |
| Total |  |  | 3537230 | 11594267 |  |  |  |  |  |

### 2. Toxicity test and efficacy of natural oil blend formulation (NOBF) in PAMs culture against African swine fever virus

The *in vitro* level trials using the natural oil blend formulation were conducted to evaluate its efficacy against African swine fever virus (ASFV). The natural oil blend had anti-viral properties and deactivated the virus (Johanna et al., 2014). The anti-viral activity of all the natural oils tested could be demonstrated for the enveloped viruses. Lavender natural oil consists primarily of monoterpenoids and sesquiterpenoids as well as linalool dominate, with moderate levels of lavandulyl acetate, terpinen-4-ol and lavandulol. 1,8-cineole and camphor are also present in low to moderate qualities. Linalol has antimicrobial, anti-inflammatory, and mood alleviating effects (Shellie et al. 2002, Johanna et al. 2014). Pine oil consists mainly of alpha-terpineol or cyclic terpene alcohols and isobornyl acetate. Pine oil is a phenolic disinfectant that is mildly antiseptic and has anti-fungal, anti-bacterial, and anti-viral properties due to isobornyl acetate (Shellie et al. 2002, Johanna et al. 2014). Eucalyptus oil has a history of full application as a pharmaceutical, antiseptic, repellent, flavoring, fragrance, and for industrial use. Eucalyptus oil has anti-bacterial, anti-viral, and anti-inflammatory effects; pre-clinical results also show that eucalyptus oil stimulates the innate cell-mediated immune response by its effects on the phagocytic ability of human monocyte-derived macrophages (Shellie et al. 2002, Johanna et al. 2014, Juergens et al. 2003 and 2004, Serafino et al. 2008 and Gobel et al. 1994). Cineole present in eucalyptus oil shows potential anti-viral activity against Herpes Virus and Yellow Fever Virus. Its activity has also been established against viral envelope structures (Jurgen et al. 2009, 2005; Schnitzler et al. 2007, Schnitzler et al., 2011; Meneses et al. 2009. The natural oils affect the viral envelope, which is necessary for adsorption or entry into host cells. In particular, monoterpenes have shown increased cell membrane fluidity and permeability, altering the order of membrane proteins (Schnitzler et al. 2010).

The toxicity of the natural oil blend formulation (NOBF) in PAMs cultures was tested in PAM cells along with the trial. PAM cell death was recorded up to dilution six at 36 hours of observation. Cell death occurred in dilution seven wells at 48 hours and dilution eight at 72 hours of culture. No PAM cells died in dilution nine and onwards until the last observation at 120 hours.

The obtained results show that the NOBF in dilutions 1 to 6 i.e., up to 0.08 ml, inhibited the growth of PAM cells (Figure 3).

**Figure 3:**
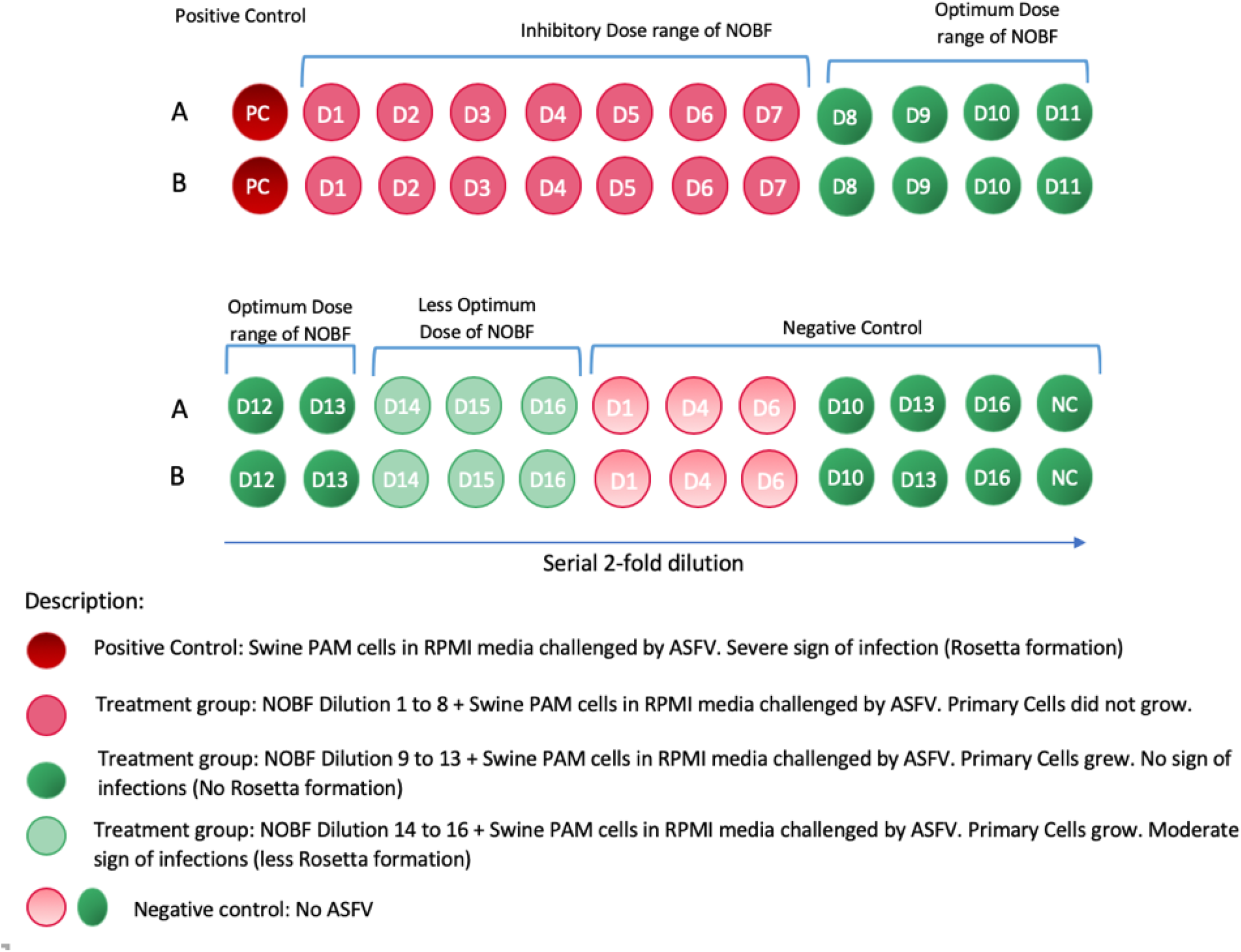
The illustration shows the result outcome of *in vitro* test. The inhibitory dose of the NOBF was observed from dilution 1 to 7. The optimum and effective dose of NOBF against ASFV was observed from dilutions 8 to 13. The less effective dose started from dilutions 14. The different dilutions of negative control (without ASFV) indicated the inhibitory and not-inhibitory doses of NOBF on PAM cells.

**Figure 4:**
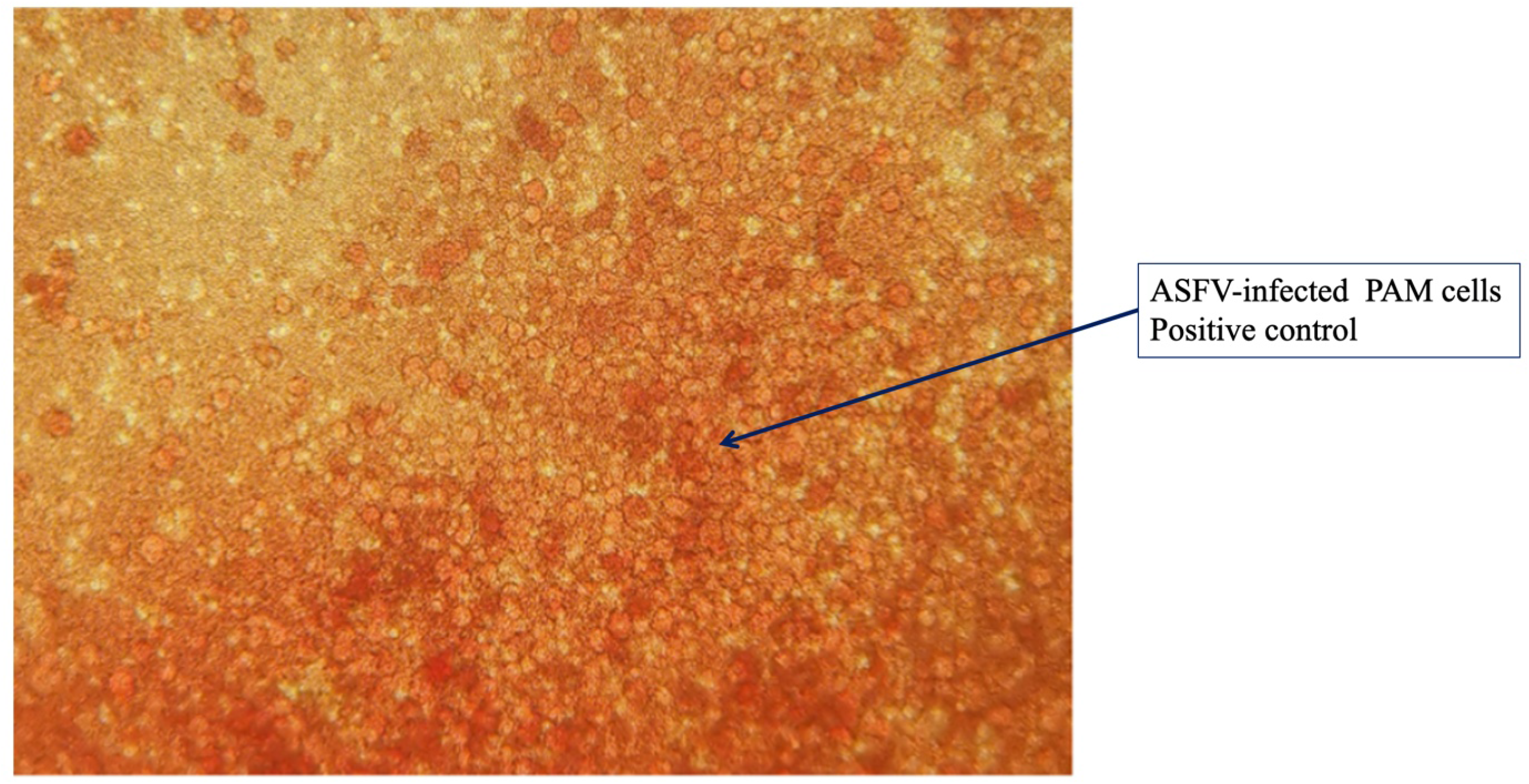
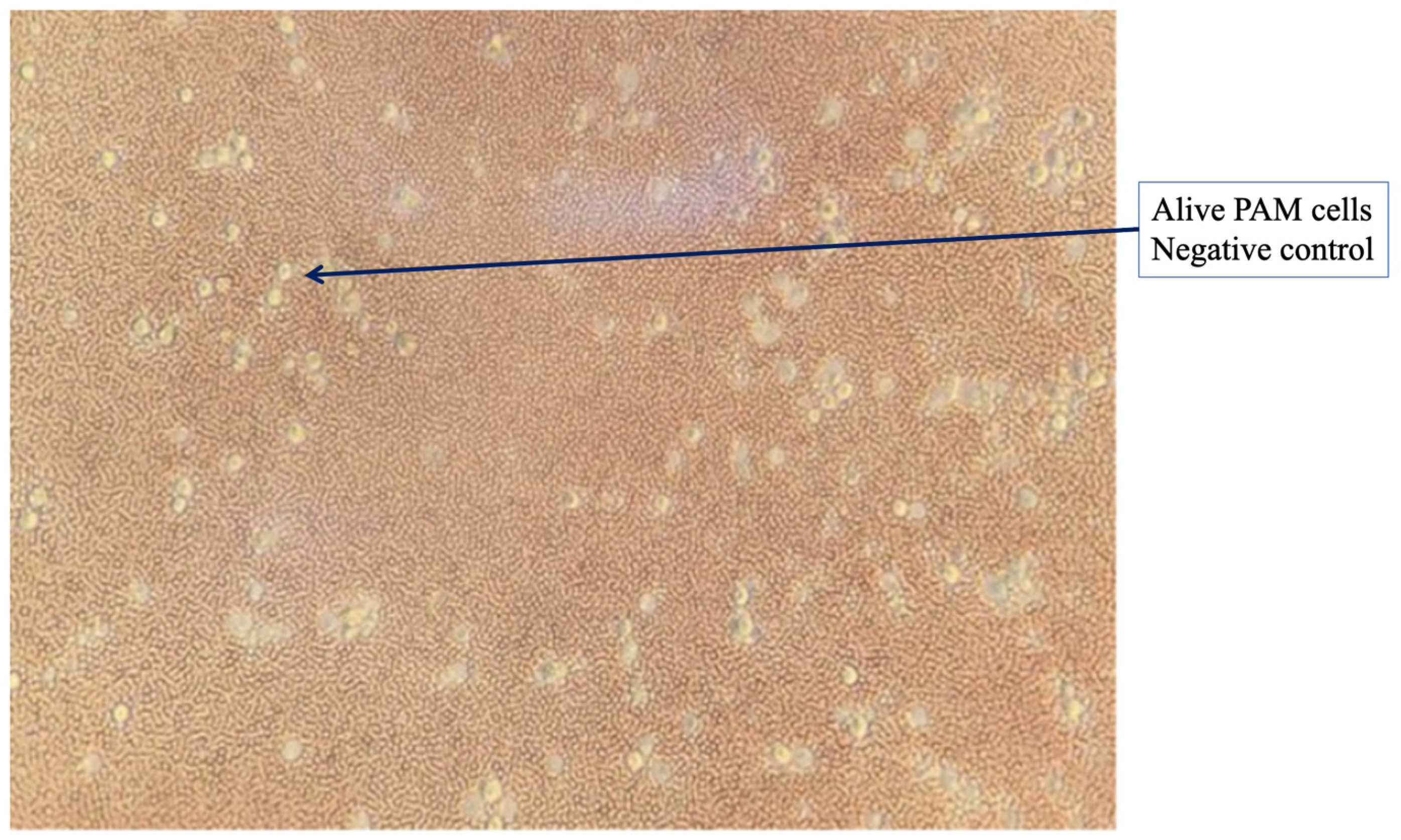
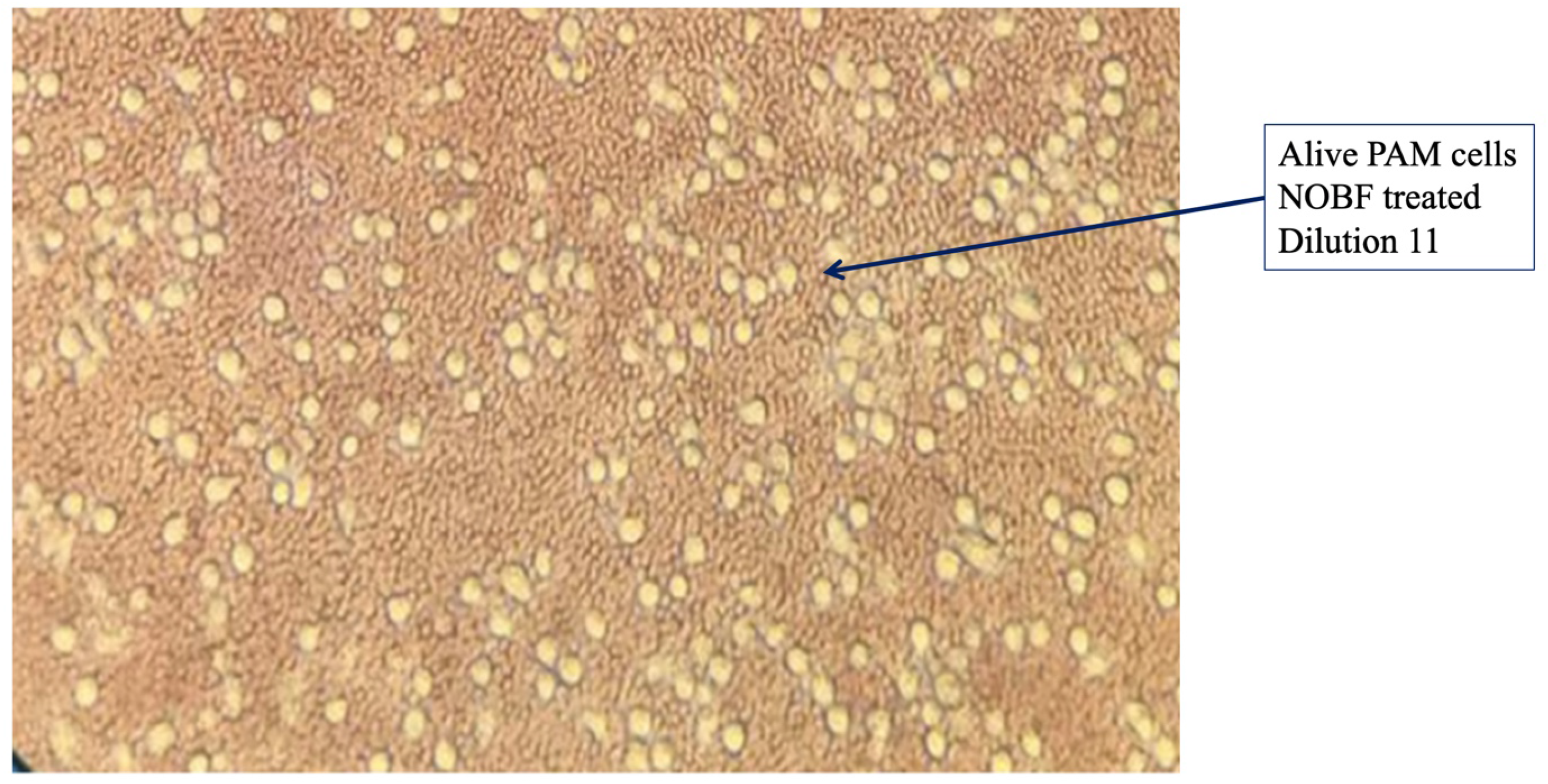
Representative microscopic images of positive control i.e. PAM cells inoculated with ASFV (a), negative control i.e. PAM cells not inoculated with ASFV and PAM cells inoculated with ASFV treated in dilution 11or 250 ppm of NOBF for 130 h (c). Images were taken on Leica DM IL LED microscope at a magnification of 200x.

The quantity log 5 of the VNUA-ASFV-L01/HN/04/19 virus strain African swine fever virus is considered the lethal dose that can kill the pigs in 7-10 days of intramuscular challenge. The evaluation of *in vitro* anti-viral activities of natural substances is based mainly on the inhibition of cytopathic effects, the reduction or inhibition of plaque formation, and the reduction in the virus yield (Jurgen et al. 2009).

The PAM cells were isolated from the pathogen-free clean piglets. The characteristic feature of the ASF virus cells is to adsorb swine erythrocytes (hemadsorption) on its surface. This property has been successfully exploited to differentiate the ASF virus from other agents that produce diseases with symptoms likely to be confused with those observed in ASF and to develop specific techniques for ASF virus titration (Galindo et al. 2000). Erythrocyte rosettes formation around infected blood swine monocytes is a characteristic feature of ASFV-infected cells (Hess and De Tray 1960, Malmquist and Hay 1960). It is considered the standard hemadsorption test for many ASFV isolates (Enjuanes et al. 1976, Galindo et al. 2000). The detailed stepwise observation at 96 hours post-infection occurred as follows:

In *vitro* observation showed that the natural oil blend formulation (NOBF) up to dilution 13 or 0.000625 ml could inhibit or degenerate ASFV at a titer of 105HAD50 in the PAMs culture (Table 1). No HAD or rosette was observed up to 130 hours post-infection (Figure 3). Whereas positive controls showed a large number of rosettes formation.

### 3. Real-time PCR analysis of *in vitro* trial

After four freeze-thaw cycles of PAMs, the supernatants were collected and applied for total DNA extraction, then used for real-time PCR. The real-time PCR results (Ct value) indicated that the virulent ASFV strain was unable to replicate or was denatured in the PAMs cultures (Table 3). No difference in Ct value was observed in the initial ASFV input control as compared to the natural oil blend group at dilutions 10, 11, and 12 after 130 hours post-infection in PAMs (Table 3). Remarkably, the difference in the Ct value of the natural oil blend group was statistically significant compared with the ASFV positive control group (Table 3).

**Table 3.** Real-time PCR quantification of ASFV replication in treatment groups *in vitro*

| No | Groups | Dilution | HAD<sup>β</sup>/real-time PCR |  |
| --- | --- | --- | --- | --- |
|  |  |  | Rosetta formation | Real-time PCR Ct value (mean) |
| 1 | Group 1: Natural oil blend formulation | 10 | No | 25.96 |
| 2 |  | 11 | No | 25.89 |
| 3 |  | 12 | No | 25.16 |
| 4 |  | 13 | No | 23.97 |
| 5 | Group 2: Positive control |  | Yes | 17.84 |
| 6 | Group 3: Negative control |  | No | NA |
| 7 | Input control of ASFV<sup>a</sup> |  |  | **25.74** |

<sup>a</sup> Initial ASFV dose that is used to apply all groups except negative control group.

## CONCLUSION

Although ASF was first described almost a century ago, controlling the disease has proven to be a challenge, mainly because no effective treatment or vaccine is available. Live-attenuated vaccines that are widely used experimentally protect some animals against infection challenges with a homologous virus, but not with a heterologous one, and most of them become carriers of the ASF virus. Inactivated vaccine or viral protein vaccines do not appear to induce any significant protection. ASF virus genes may play a role in the modulation of protection that has been researched. However, until now, the only treatment for ASF is eradication, based on the control of animal transport and vectors and the early detection and slaughter of infected and carrier animals. The role of natural oil extracts in virus eradication has been recognized and considered. In this study, we try to demonstrate that the natural oil blend formulation can degenerate the African Swine Fever Virus (ASFV).

The trial outcome can be summarized in the following way: the natural oil blend formulation (NOBF) application deactivated the ASF virus at the lowest concentration of 0.000635 ml. As a continuation of the study, the next step is to conduct *in vivo* trials to optimize the dose and delivery route of the natural oil blend formulation (NOBF) and to establish it as an anti-ASFV candidate.

## ACKNOWLEDGEMENTS

We would also like to show our gratitude to CeRAF, Vietnam, VNUA, Vietnam, and Asclepius Pharmaceutical Sciences, Singapore, and PT. Central Proteina Prima Tbk., Indonesia, and the Ministry of Science and Technology for support in making this research possible.

